# Local genetic correlation gives insights into the shared genetic architecture of complex traits

**DOI:** 10.1101/092668

**Authors:** Huwenbo Shi, Nicholas Mancuso, Sarah Spendlove, Bogdan Pasaniuc

## Abstract

Although genetic correlations between complex traits provide valuable insights into epidemiological and etiological studies, a precise quantification of which genomic regions contribute to the genome-wide genetic correlation is currently lacking. Here, we introduce *ρ*-HESS, a technique to quantify the correlation between pairs of traits due to genetic variation at a small region in the genome. Our approach only requires GWAS summary data and makes no distributional assumption on the causal variant effects sizes while accounting for linkage disequilibrium (LD) and overlapping GWAS samples. We analyzed large-scale GWAS summary data across 35 complex traits, and identified 27 genomic regions that contribute significantly to the genetic correlation among these traits. Notably, we find 7 genomic regions that contribute to the genetic correlation of 12 pairs of traits that show negligible genome-wide correlation, further showcasing the power of local genetic correlation analyses. Finally, we leverage the distribution of local genetic correlations across the genome to assign putative direction of causality for 15 pairs of traits.

## Introduction

Genomic regions that harbor variants contributing to multiple traits provide valuable insights into the underlying biological mechanisms with which genetic variation impacts complex traits^1,2,3,4,5,6,7^. Therefore, both the identification of new such regions as well as quantifying the correlation in causal effects at known shared regions, is of great importance in epidemiological and etiological studies. For example, genetic variants associated with multiple traits in genome-wide associations studies (GWAS), can be used as instrumental variables in Mendelian randomization analyses to identify causal relationships among complex traits^7,8,9,10^. Unfortunately many risk variants are left undetected by existing GWAS due to a combination of high polygenicity (i.e. many variants of small effects) and sample sizes which limits the power to detect genetic variants of small effect^11^. To improve accuracy at sub-GWAS significant loci, recent works^1,2^ proposed to utilize the posterior probability of two traits sharing a causal variant at each locus to detect genetic overlap. Although powerful in detecting shared genetic risk variants, the posterior probability does not convey the direction or magnitude of the genetic effect at the overlapped loci^1,2^. Alternative approaches have used genetic correlation (i.e. the correlation between the causal effects), that summarizes both direction and magnitude of effects, to gain insights into genetic overlap of complex traits^12,13^. Due to polygenicity assumptions, genetic correlation has been only investigated in genome-wide context by aggregating information across all variants in the genome^14,15^. In this work, we investigate local genetic correlation (i.e. correlation between a pair of traits only due to genetic variants from a small region in the genome) as means to dissect the genetic sharing between pairs of traits.

Traditional methods to estimate genetic correlation between a pair of traits rely on pedigree or family data, requiring phenotype measurements of the traits on the same set of individuals^13,14^. While more recent bivariate-REML^15^ and HE-regression^16^ methods provide the convenience to estimate genetic correlation from genotype and phenotype data of unrelated individuals, they are hindered by the lack of availability of large-scale individual-level data due to privacy concerns^12,14,17,18^. A recently proposed method, cross-trait LD score regression (LDSC) circumvented this hindrance by estimating genetic correlation from GWAS summary data, which are publicly available from most large GWAS consortia^19,20^, and created an atlas of genetic correlation across multiple human complex traits and diseases^14^. However, due to model assumptions on trait polygenicity, cross-trait LDSC has been only applied for estimating genome-wide genetic correlation^12,14^, which quantifies the direction and magnitude of genetic sharing across the entire genome^14^.

In this work we introduce *ρ*-HESS, a method to estimate the local genetic correlation between a pair of traits at each region in the genome from GWAS summary data, while accounting for overlapping GWAS samples and linkage disequilibrium (LD). Our method estimates the contribution to the genetic correlation of typed variants for each LD block in the genome; we utilize approximately independent LD blocks of roughly 1.5Mb in size on the average^21^. We make no distributional assumption on the causal effect sizes by treating them as fixed quantities, which allows for a broad range of causal genetic architectures at small regions in the genome. Our approach can be viewed as a natural extension to pairs of traits of recently proposed methods that quantify local SNP-heritability from GWAS summary data under a fixed effects model^18^. Through extensive simulations, we demonstrate that given in-sample LD, *ρ*-HESS yields unbiased estimates of local genetic covariance, and approximately unbiased estimates of local genetic covariance and correlation when LD is estimated from external population reference panels such as the 1000 Genomes data^22^.

We analyzed GWAS summary data across 35 complex traits and identified 234 pairs of traits with significant genome-wide genetic correlation; these include previously reported correlations (e.g., lipid traits) as well as correlations between traits not reported before (e.g., blood related traits). Second, we identify 27 genomic regions that show significant local genetic covariance as well as local SNP-heritability across 29 pairs of traits. For example, *ρ*-HESS estimates a local genetic correlation of −0.95 (s.e. 0.16) for the region chr2:21-23M, which harbors the APOB gene for HDL and TG. Notably, 7 (out of the 27) significantly correlated genomic regions across 12 pairs of traits are significant in local analyses, although the genome-wide genetic correlation is not significantly different from 0. For example, when considering genetic correlation between mean cell volume (MCV) and platelet count (PLT), region chr6:134-136M shows a local genetic correlation of 1.0 (s.e. 0.16), whereas the genome-wide genetic correlation is negligible (0.01 s.e. 0.04). This shows that these traits harbor genetic overlap at a local level (e.g., due to pleiotropy and/or shared pathways) and emphasize the power of local correlation analysis.

Having estimates of local genetic correlation at every region in the genome allows us to perform bi-directional analyses to assign putative direction of causality for a pair of traits. Intuitively, all loci that harbor variants that contribute to one trait will have a consistent direction of effect on the second trait under a causal model, whereas under no causal model we expect no consistency in the direction of genetic effects (see Figure 1). This extends previously proposed works^2^ from individual variant effects to local genetic covariances. We assign putative direction of causality for 15 pairs of traits. Reassuringly, our analyses show that height causally decreases body mass index (BMI), and that BMI causally increases triglyceride (TG), consistent with expectation and previous works^2,3^. We also identify putative causality between previously unreported traits. For example, our analyses suggest that schizophrenia (SCZ) causally increases the risk for ulcerative colitis (UC), providing possible explanation to why prevalence of UC is higher in SCZ patients but not vice versa^23^. Interestingly we also find evidence of age at menarche (AM) causally decreases the level of triglyceride (TG). Overall, our results motivate further work in confirming putative causality direction among these traits.

**Figure 1:**
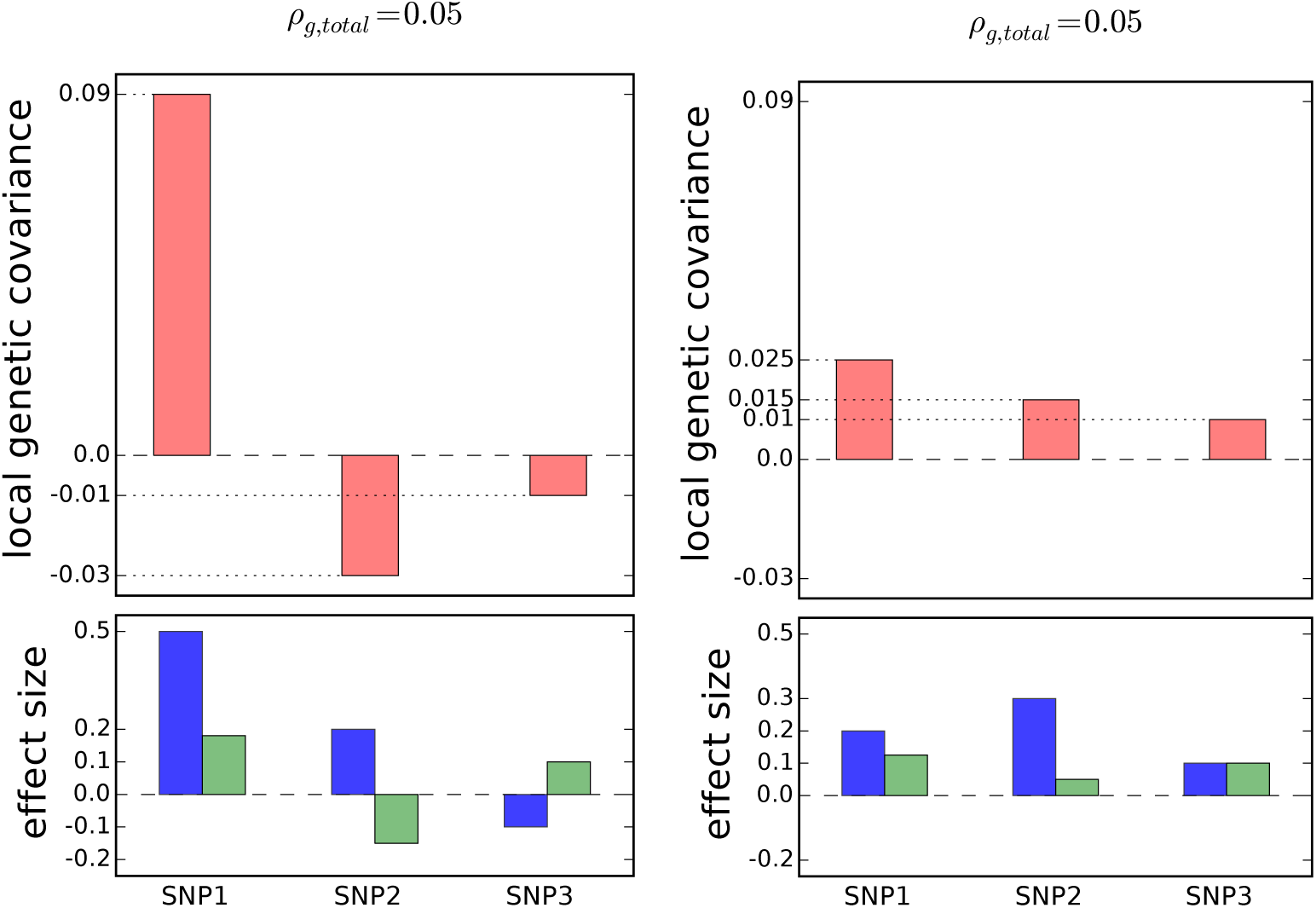
Examples of two different distributions of local genetic covariances (with effect sizes shown at the bottom) that result in the same total genetic covariance (*ρ_g,total_* = 0.05). In the left example, the total genetic covariance is a summation of a large positive local genetic covariance at SNP1 and two smaller negative local genetic covariances at SNP2 and SNP3 (e.g, SNPs 2 and 3 impact traits through a different pathway than SNP1). In the right side the total genetic covariance is a summation of small positive local genetic covariances (e.g., all three SNPs impact both traits through the same pathway). Positive local genetic covariance can be interpreted as a locus driving a pathway that regulates two traits in the same direction, and negative local genetic covariance the opposite direction.

## Results

### Overview of methods

Genetic correlation measures the similarity between a pair of traits driven by genetic variation (e.g., correlation on causal effect sizes), and enjoys wide applications in understanding relations between complex traits^12,24,25^. Genetic correlation is traditionally estimated as a single measure across the entire genome to capture the contribution of genetic variation across the entire genome to the correlation between phenotypes. Here, we introduce local genetic correlation, the similarity between pairs of traits driven by genetic variation localized at a specific region in the genome (e.g., one LD block), as a principled way to dissect the genome-wide genetic correlation between traits. For example, a high genome-wide genetic covariance can be driven by one or a few genomic regions containing a shared risk variant, or by a large number of regions each with a small contribution (see Figure 1). The distribution of local genetic covariances reflects causality relations (where all risk variants for one trait are risk variants for the other trait) and/or pleiotropic regions (risk variants contributing to both traits through different pathways).

Under the assumption that true causal effects of genetic variants on the two traits are known, the local genetic covariance between the two traits is simply *β*^T^***V****γ*, where *β* and *γ* are the causal effect vectors and *V* is the pairwise correlation between variants (i.e. linkage disequilibrium, LD) (see Methods). A traditional GWAS estimates marginal effect sizes at each variant that contain statistical noise due to finite sample size and are confounded by LD^19^. We deconvolute LD and account for the statistical noise to estimate the local genetic covariance as:

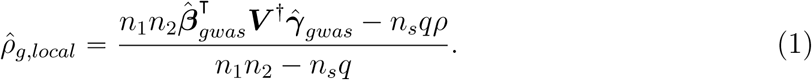

Here, 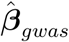 and 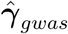 are the vectors of standardized marginal GWAS effect sizes, *n_1_* and *n_2_* the sample size of each GWAS study, *n_s_* the number of samples shared across the two GWASs, *q* the rank of the LD matrix ***V***, *ρ* the total phenotypic correlation between the two traits and ***V***^†^ is Moore-Penrose pseudoinverse of the LD matrix (***V***). The variance of the estimator follows from bilinear form theory, and has magnitude approximately on the order of 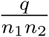 (see Methods). To obtain local genetic correlation, we standardize the covariance as 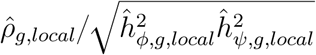 where 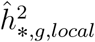 are the local SNP-heritability^18^ estimates for the two traits.

When in-sample LD matrix (***V***) is available, Equation (1) yields an unbiased estimator of local genetic covariance (see Results). When individual-level genotype data is not available, we use LD matrix (***V̂***) estimated from external reference panels (e.g., 1000 Genomes Project). The small sample size of external reference panels creates statistical noise in the estimated external reference LD matrix ***V̂*** and results in biased estimates of local genetic covariance and correlation. To account for the statistical noise, we apply truncated-SVD regularization (see Methods).

### Accuracy of local correlation estimation in simulations

We evaluated the performance of *ρ*-HESS through extensive simulations across various disease architectures. First, we assessed the performance of our approach across varying simulated local genetic covariances. As expected, when in-sample LD matrix is available, *ρ*-HESS provides an unbiased estimate of local genetic covariance (see Figure 2). For completeness, we also adapted cross-trait LDSC^14^ for local estimation and observed biased results; this is expected as cross-trait LDSC is not designed for local estimation under fixed effects at causal variants. For example, for simulated local genetic covariance of 1.7 × 10^−4^, *ρ*-HESS yields an estimate of 2.0 × 10^−4^ (s.e. 4.3 × 10^−5^), whereas cross-trait LDSC yields an estimate of 6.0 × 10^−4^(s.e. 3.4 y 10^−5^). Second, we assessed the performance of local genetic correlation (i.e. the local genetic covariance divided by the square root of their respective local heritability estimates). Overall, the local genetic correlation estimates are approximately unbiased; e.g., for a simulated local genetic correlation of 0.35 *ρ*-HESS yielded an estimate of 0.22 (s.e. 0.19) (see Figure 2). Third, we quantified the performance of our approach in the case when in-sample LD is unavailable and needs to be estimated from external reference panels. We observe that the *ρ*;-HESS estimates of local genetic covariance and correlation are still approximately unbiased (see Figure 2) when truncated-SVD is used for regularizing the LD matrix (see Methods). Noticeably, the standard errors of ρ-HESS estimates also decrease due to the truncated-SVD regularization. Finally, we compared the performance of local genetic correlation estimation in simulations with different degrees of polygenecity. Overall, we observe that *ρ*-HESS is not sensitive to the underlying polygenicity of the trait, and provides nearly unbiased and consistent estimates of local genetic covariance and correlation (see Figure 2).

**Figure 2:**
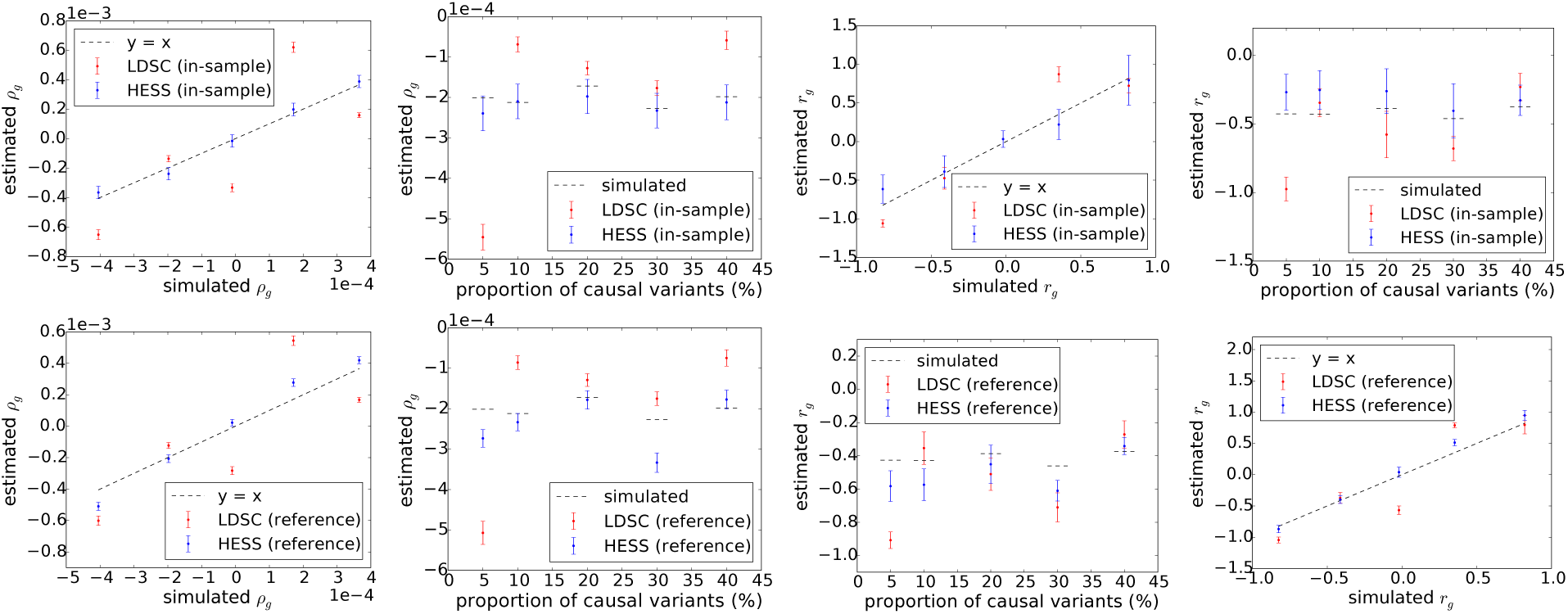
*ρ*-HESS provides unbiased estimates of covariance when in-sample LD is available (top left) and nearly unbiased estimates of correlation/covariance when LD is estimated from a reference panels (bottom and top right) Mean and standard errors are computed based on 500 simulations. Error bars represent 1.96 times the standard error on each side.

### Partitioning the genetic correlation across local genomic regions

We analyzed GWAS summary data of 35 complex traits to obtain local genetic covariances/correlations at 1,703 approximately LD-independent regions in the genome (˜1.5Mb on the average)^21^. First, we aggregated the local estimates into genome-wide estimates of genetic correlation (see Methods, Supplementary Figure 2–8) and found a high degree of concordance with genetic correlation estimated by cross-trait LDSC regression (*R* = 0.74, see Figure 3, Supplementary Figure 1). We note that our estimator provides consistently lower estimates than LDSC for pairs of traits from the same consortium where our approach assumes full sample overlap and therefore is more conservative (see Discussion). We report 234 pairs of traits with significant genome-wide genetic correlation after correcting for 595 pairs investigated (*p* < 0.05/595). These include previously reported genetic correlations, e.g. body mass index (BMI) and triglyceride (TG), as well as complex traits that have not been studied before using genetic correlation, e.g. type 2 diabetes (T2D) and ulcerative colitis (UC).

**Figure 3:**
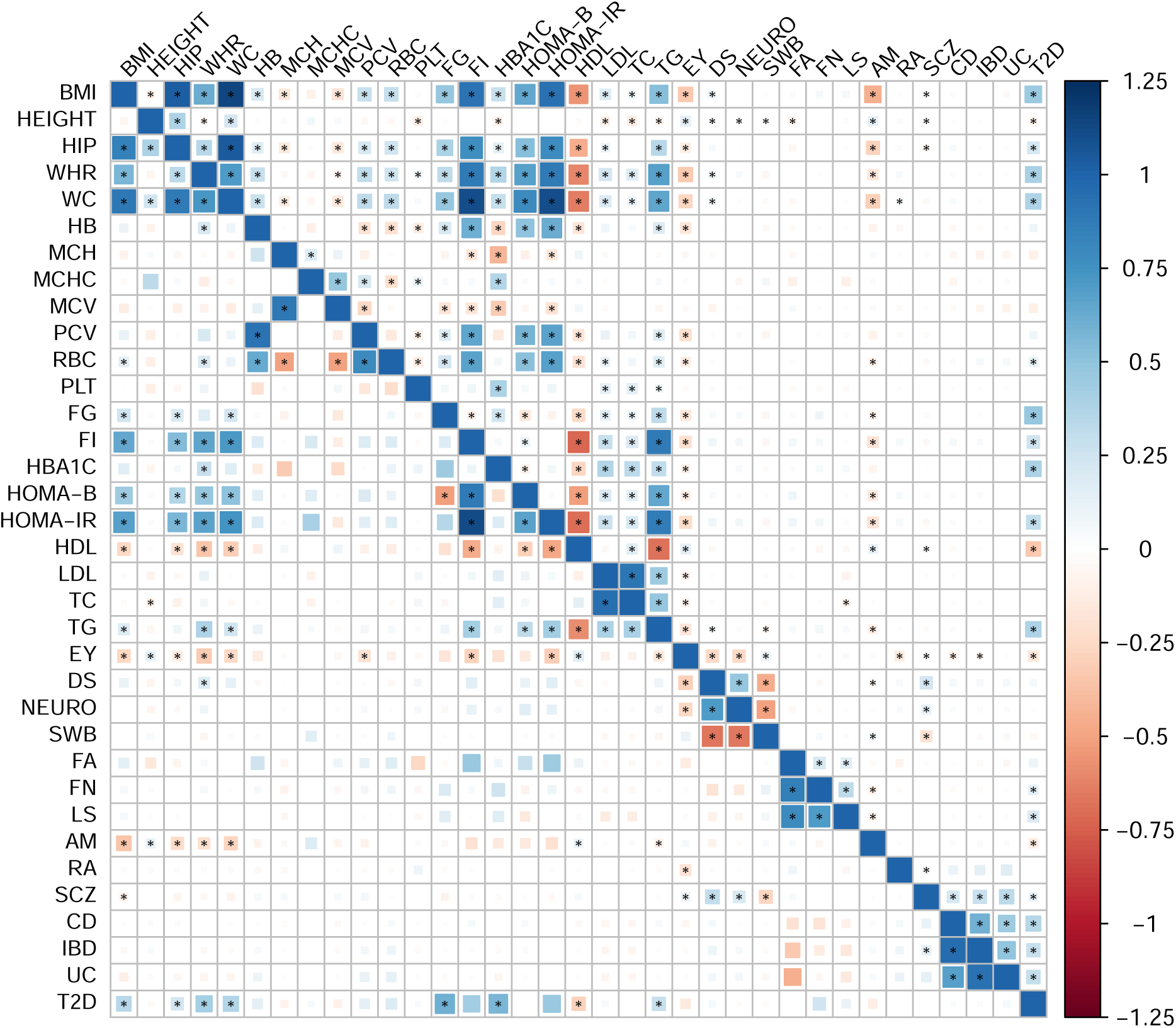
Genetic correlation across the 35 complex traits obtained by *ρ*-HESS (top half) and cross-trait LDSC^1414^ (bottom half). The magnitude of the correlation is represented by the color and the size of the square. Among the 595 pairs of traits, *ρ*-HESS (LDSC) identified 234 (99) pairs showing significant genetic correlation (marked with dots)

Next, we searched for genomic regions that disproportionately contribute to the genetic correlation of the 35 analyzed traits; we excluded the HLA region due to complex LD patterns. We identify 27 genomic regions that show significant local genetic correlation (two-tailed *p* < 0.05/1703/595) as well as significant local SNP-heritability (one-tailed *p* < 0.05/1703/35) (see Figures 4,5, Supplementary Figures 9, Supplementary Table 1 and Table 2). 22 out of 27 regions contain GWAS significant SNPs for both traits, whereas 5 of these loci contain GWAS significant hits for only one or none of traits and can be viewed as new risk regions for these traits. For example, the estimate of local genetic correlation between HDL and TG at chr11:116-117Mb is −0.82 (s.e. 0.07), suggesting highly shared genetic architecture at this locus for HDL and TG. Indeed, the locus chr11:116-117M harbors the APOA gene, which is known to be associated with multiple lipid traits^26^.

Since genetic correlation is an aggregated manifestation of local genetic covariance, for pairs of traits with highly positive or negative genetic correlation, we expect the distribution of local genetic covariances to be shifted towards the positive or negative side; whereas for pairs of traits with low genetic correlation, we expect the distribution of local genetic covariances to be centered around zero (see Figure 6,7,8). Indeed, pairs of traits with higher genome-wide genetic correlation tend to harbor more loci with significant local genetic covariance (see Figure 4). For instance, only one locus exhibits significant local genetic covariance for the pair of traits age at menarche (AM) and height (*r_g_* = 0.13, s.e. 0.01), whereas four loci show significant local genetic covariance for the pair of traits LDL and TG (*r_g_* = 0.44, s.e. 0.02).

**Figure 4:**
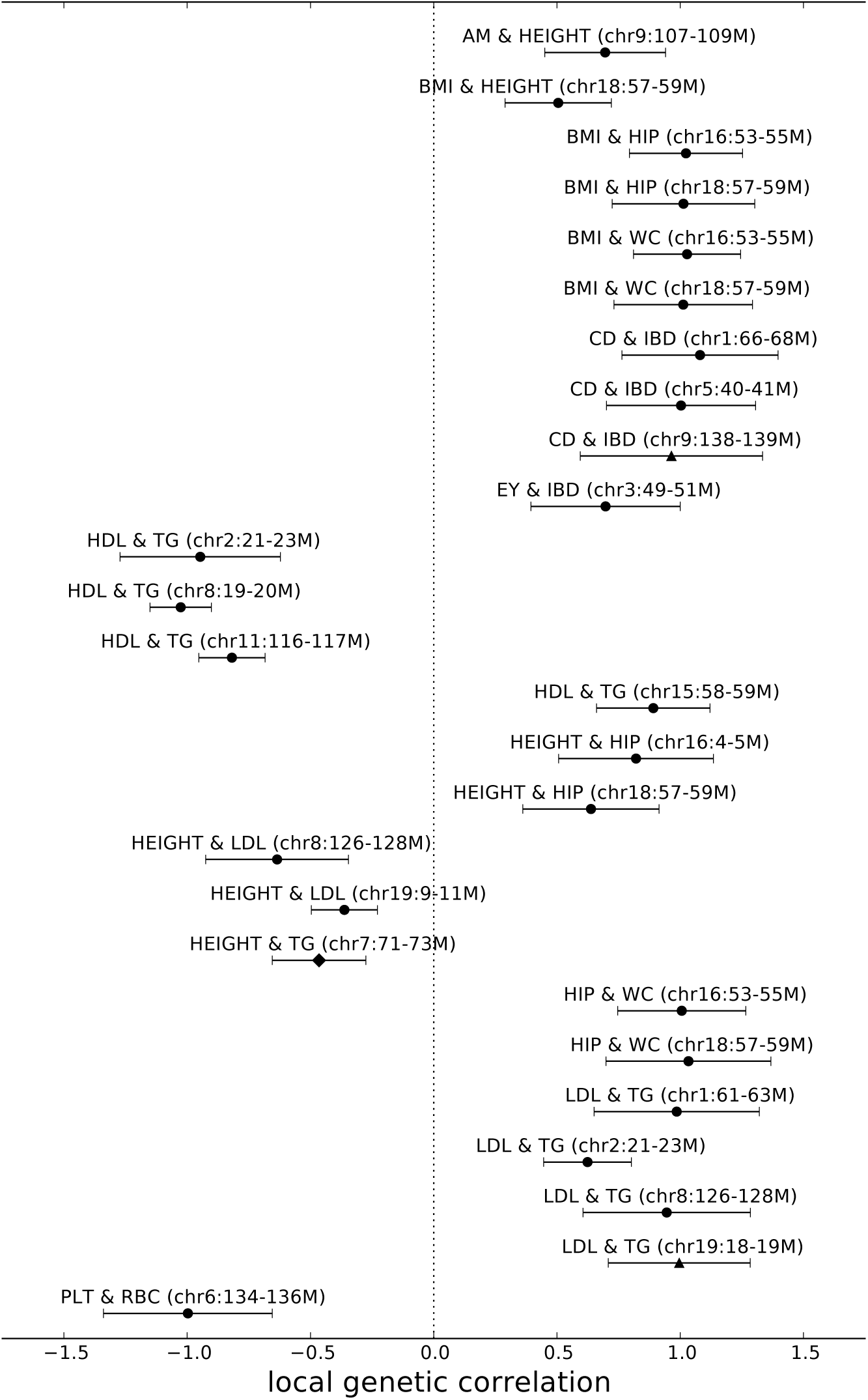
Local genetic correlation at loci displaying significant local genetic covariance and SNP-heritability for pairs of traits with significant genome-wide genetic correlation. To simplify presentation, we excluded all pairs of traits involving TC (see Supplementary Figure 9). We obtain standard error estimates through parametric bootstrapping. Error bars represent 1.96 times the standard error on both sides. Here, triangle represents loci that lack GWAS risk variant for both traits; diamond represents loci that harbor GWAS risk variants for one of the traits; and circle represents loci that contain GWAS risk variants for both traits.

**Figure 6:**
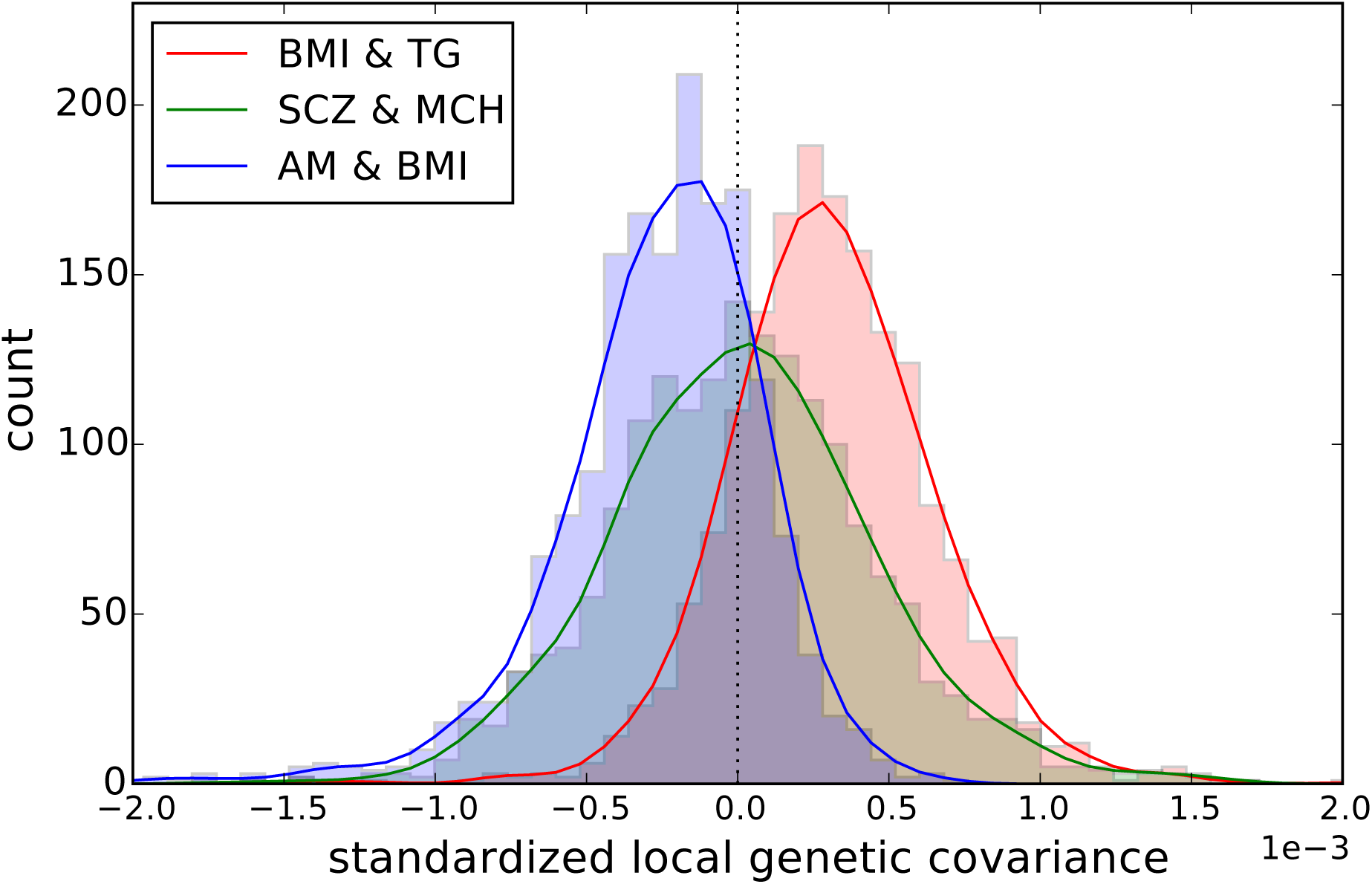
Distribution of standardized local genetic covariance (local genetic covariance standardized by the square roots of heritability of two traits) for the pairs of traits BMI and TG, SCZ and MCH, AM and BMI. Pairs of traits with positive (negative) genome-wide genetic correlation show a shift in the distribution of standardized local genetic covariance away from 0.

### Local correlations for pairs of traits with negligible genome-wide correlation

Although many complex traits are known to share risk loci, some pairs of traits show negligible genome-wide genetic correlation. For example HDL and LDL share several GWAS risk loci^26^ but the genome-wide genetic correlation is negligible (−0.05, s.e. 0.02)^14^(see Figure 3). The absence of significant genome-wide genetic correlation between these pairs of traits can be attributed to either symmetric distribution of local genetic covariance (positive local genetic covariance cancels out negative local genetic covariance, see Figure 1) and/or lack of power to declare significance for genome-wide genetic correlation. Thus, we hypothesize that at the locus-specific level, many loci may manifest significant local genetic covariance even if the genome-wide genetic correlation between a pair of traits is not significant. Indeed, 9 genomic regions show significant local genetic correlation (two-tailed *p* < 0.05/1703) for HDL and LDL (see Figure 7). Some of these loci, e.g. chr2:21M-23M, chr11:116M-117M, and chr19:44M-46M, harbor genes (APOB, APOA, and APOE, respectively) that are well known to be involved in lipid genetics^26,27,28^. Across all pairs of traits with non-significant genome-wide correlation, we identify 7 regions across 12 pairs of traits with significant local genetic covariance (two-tailed *p* < 0.05/1703/595) as well as significant local SNP-heritability (one-tailed *p* < 0.05/1703/35) (see Figure 5, Supplementary Table 2). Most of these loci also show significant local genetic correlation, suggesting high similarity in the effects of genetics on traits at these loci (see Figure 5, Supplementary Table 2). For example the region chr6:134-136M harbors the blood-trait-associated gene HBS1L^29,30^, and contributes to local genetic covariance across many blood traits (MCH, MCV, RBC, and PLT).

**Figure 5:**
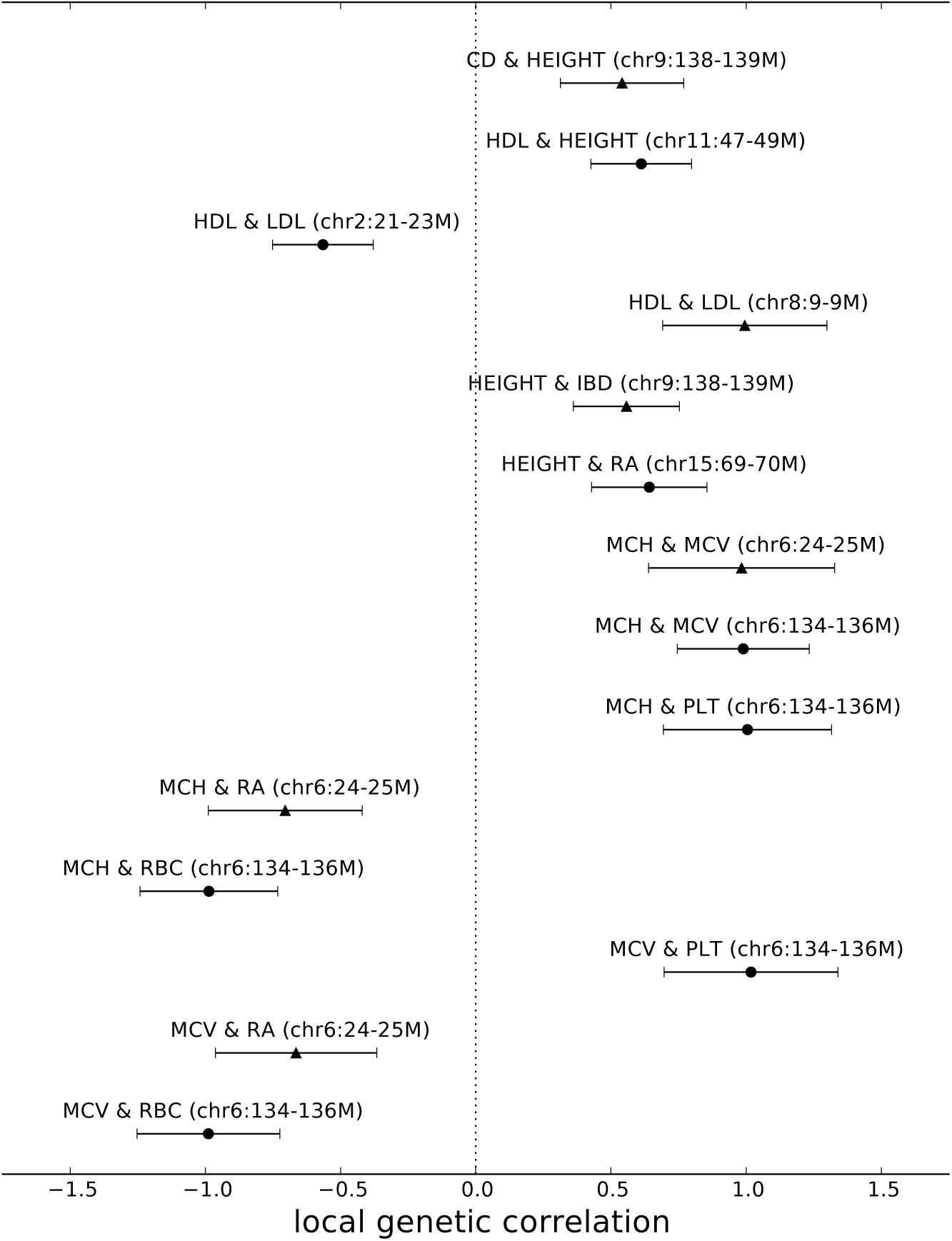
Local genetic correlation at loci displaying significant local genetic covariance and SNP-heritability for pairs of traits without significant genome-wide genetic correlation. We obtain standard error estimates through parametric bootstrapping. Error bars represent 1.96 times the standard error on both sides. Here, triangle represents loci that lack GWAS risk variant for both traits; diamond represents loci that harbor GWAS risk variants for one of the traits; and circle represents loci that contain GWAS risk variants for both traits.

**Figure 7:**
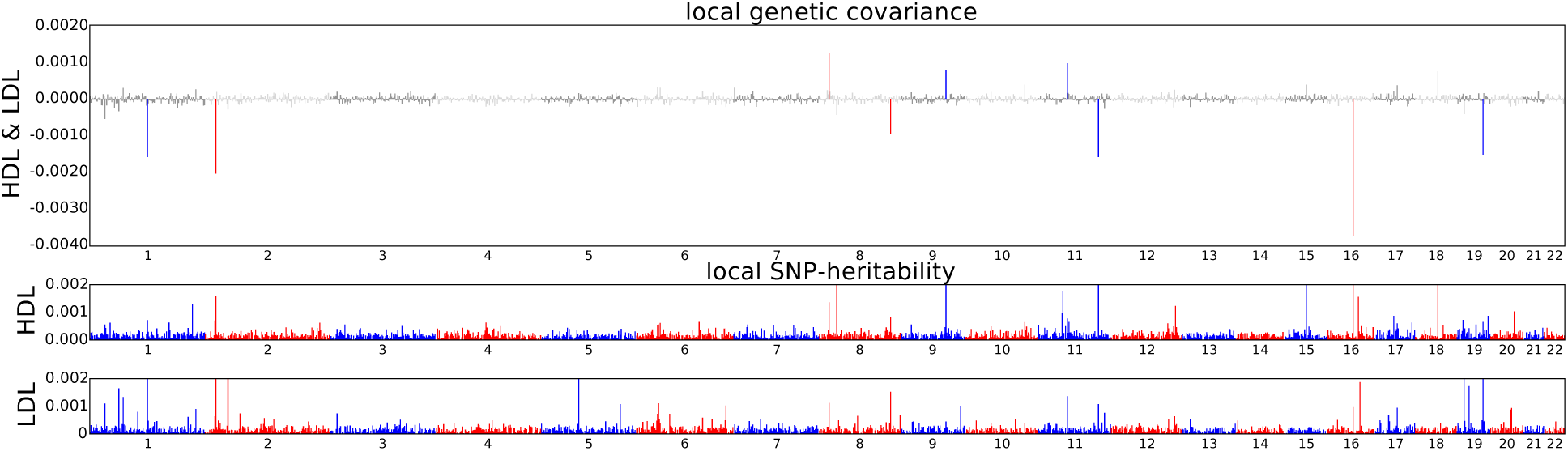
Manhattan-style plots showing the estimates of local genetic covariance for the pairs of traits HDL and LDL. Although the genome-wide genetic correlation between HDL and LDL does not reach the significance level (*p* < 0.05/595), 9 loci exhibit significant local genetic covariance.

**Figure 8:**
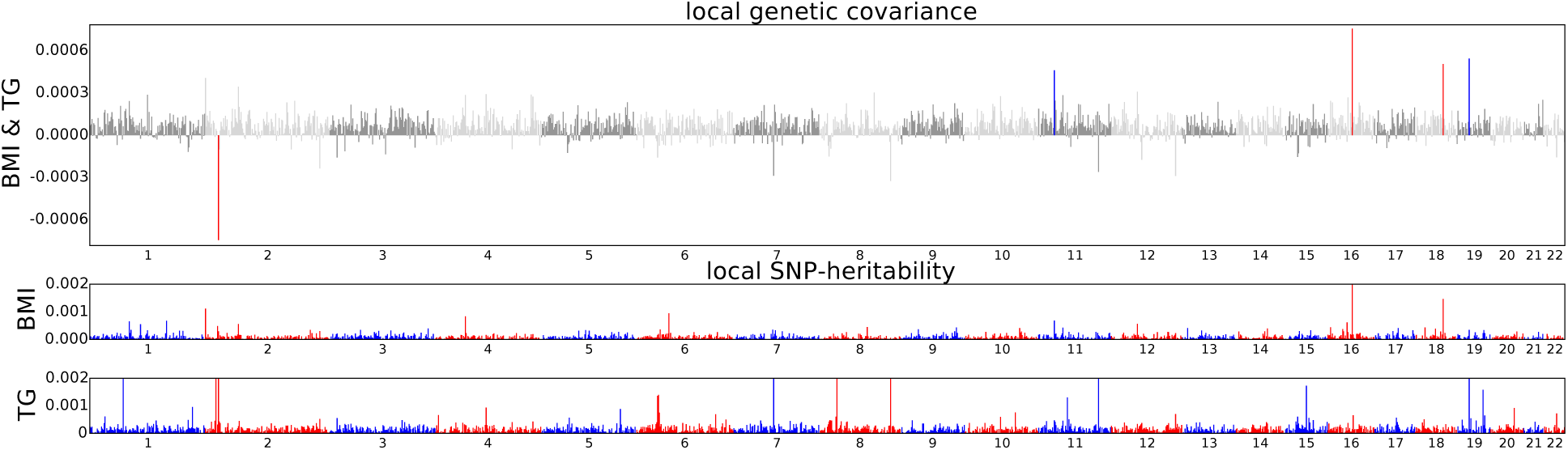
Manhattan-style plots showing the estimates of local genetic covariance for the pairs of traits BMI and TG. That the local genetic covariance between BMI and TG is mostly one-sided implies plausible causal relationship between the two traits.

### Putative causal relationships between complex traits

The distribution of local genetic correlations across the genome can be leveraged to infer putative direction of causality across pairs of traits. We illustrate this approach through 4 examples of putative causality directions (see Figure 9). First, consider the example of BMI and TG, genetically correlated genome-wide (*r̂_g_*=0.53, s.e. 0.02). The local genetic correlation restricted only to loci that harbor a GWAS hit for BMI (and no GWAS hit for TG) is significant *r̂_g,local,BMI_*=0.52, s.e. 0.05 and consistent with genome-wide correlation. In contrast, the local genetic correlation estimated at loci that harbor GWAS hits for TG (and no GWAS hits for BMI) is not significantly different from 0 (*r̂_g,local,TG_*=−0.020, s.e. 0.053). This shows that genetic variants that contribute to BMI have a consistent effect on TG whereas genetic variants associated to TG do not have a consistent effect on BMI. These two observations provide evidence favoring the model in which BMI causally increases TG (see Figure 9). Second, the pattern where one of the conditional correlations is not significant needs not be just in the positive direction. For example, BMI and height are genome-wide genetically correlated at (*r* = −0.08, s.e. 0.01), whereas *r̂_g,local,height_* is significantly negative (−0.15, s.e. 0.02), and *r̂_g,local,height_* close to zero (−0.01, s.e. 0.04). This supports a model where height causally decreases BMI (as expected, see Figure 9). We note, however, that local genetic correlations between can also be induced by shared biological pathways driven by genetic variations at these loci and/or an unobserved confounder; for example, shared genetic variants impacting a sub-phenotype that is pleiotropic for both traits could induce local genetic correlations consistent with a causal model. Thus, we exercise caution when utilizing the local genetic correlations to identify putative causality. Finally, the conditional correlation approach cannot distinguish between putative causality (or other complex model that includes unobserved phenotypes) when both conditional correlations are significant or when both correlations are not significantly different from 0 (e.g., AM-BMI and MCH-SCZ, see Figure 9)

**Figure 9:**
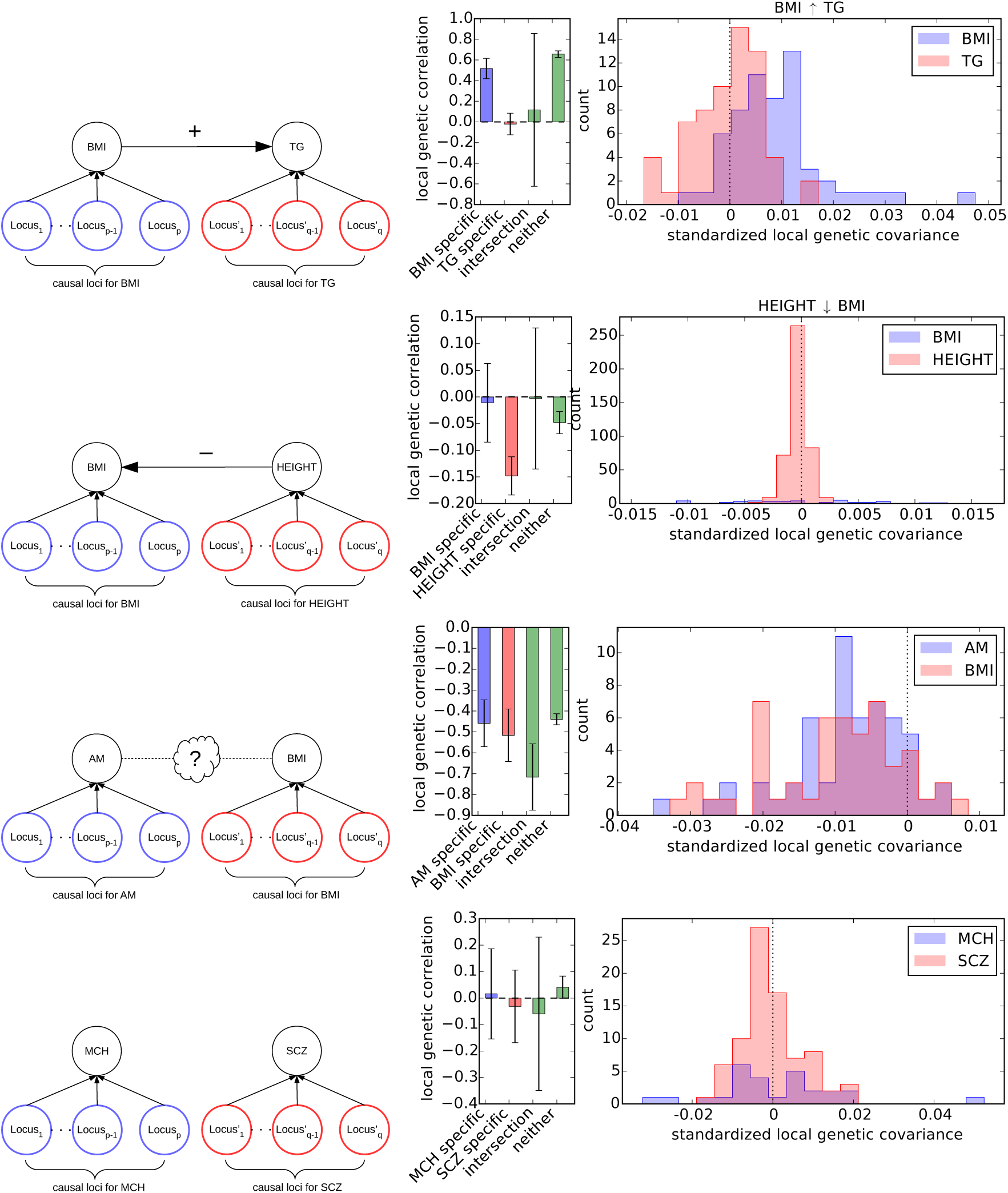
Putative causal relationships between pairs of traits. Example 1: BMI causally increases TG. Local genetic correlation estimate ascertained for loci harboring BMI risk variants is significantly greater than 0. Example 2: Height causally decreases BMI. Local genetic correlation estimate ascertained for loci harboring height risk variants is significantly less than 0. Example 3: More complicated relationships exist between AM and BMI. Local genetic correlation estimates ascertained for loci harboring risk variants for both traits are significantly less than 0. Example 4: Evidence does not support a model in which there is any causal relationship between MCH and schizophrenia (SCZ).

Through our bi-directional analyses (see Methods), we identified a total of 15 pairs of traits that support a causal directional effect (see Figure 10, Supplementary Figure 10-13). As an example, our bi-direction analyses provide evidence favoring the model, in which age at menarche (AM) causally decreases triglyceride (TG) level (*r̂_g,local,AM_* = ‒0.28 s.e. 0.06, *r̂ _g,local,TG_* = 0.04 s.e. 0.05), suggesting that the association observed between AM and TG is likely induced by the effect of AM on TG ^31,32^. Previous work also shows that association exists between schizophrenia (SCZ) and ulcerative colitis (UC), however, the direction of causality is uncertain^23^. Our bi-directional analyses favors a model in which SCZ causally increases the risk of UC (*r̂_g,local,SCZ_* = 0.30 s.e. 0.05, *r̂_g,local,UC_* = ‒0.02 s.e. 0.07). This result may provide insights into the previous observations of no increase in prevalence of SCZ in UC patients^33^. However, our result does not rule out other causal models, for example, it could be that shared genetic variants drive an immune system phenotype that is pleiotropic for both SCZ and UC, inducing the relationship between *r̂_g,local,SCZ_* and *r̂_g,local,UC_*^34,35^. Interestingly, we also observe that our bi-directional analyses support a model in which years of education (EY) causally decreases hemoglobin level (HB), LDL, TG, and the risk for rheumatoid arthritis (RA) (see Figure 10). We note that these results are consistent with previous conclusions on the causal effect of education on health^36,37^. However, we emphasize that education attainment (or other studied traits) may be confounded by other factors such as social status and that one should exercise caution when interpreting these results. Finally, our bi-directional analyses over local genetic correlation also suggest that BMI likely causally increases triglyceride (TG) and that height causally decreases BMI, consistent with our expectations and previous findings^2,3^.

**Figure 10:**
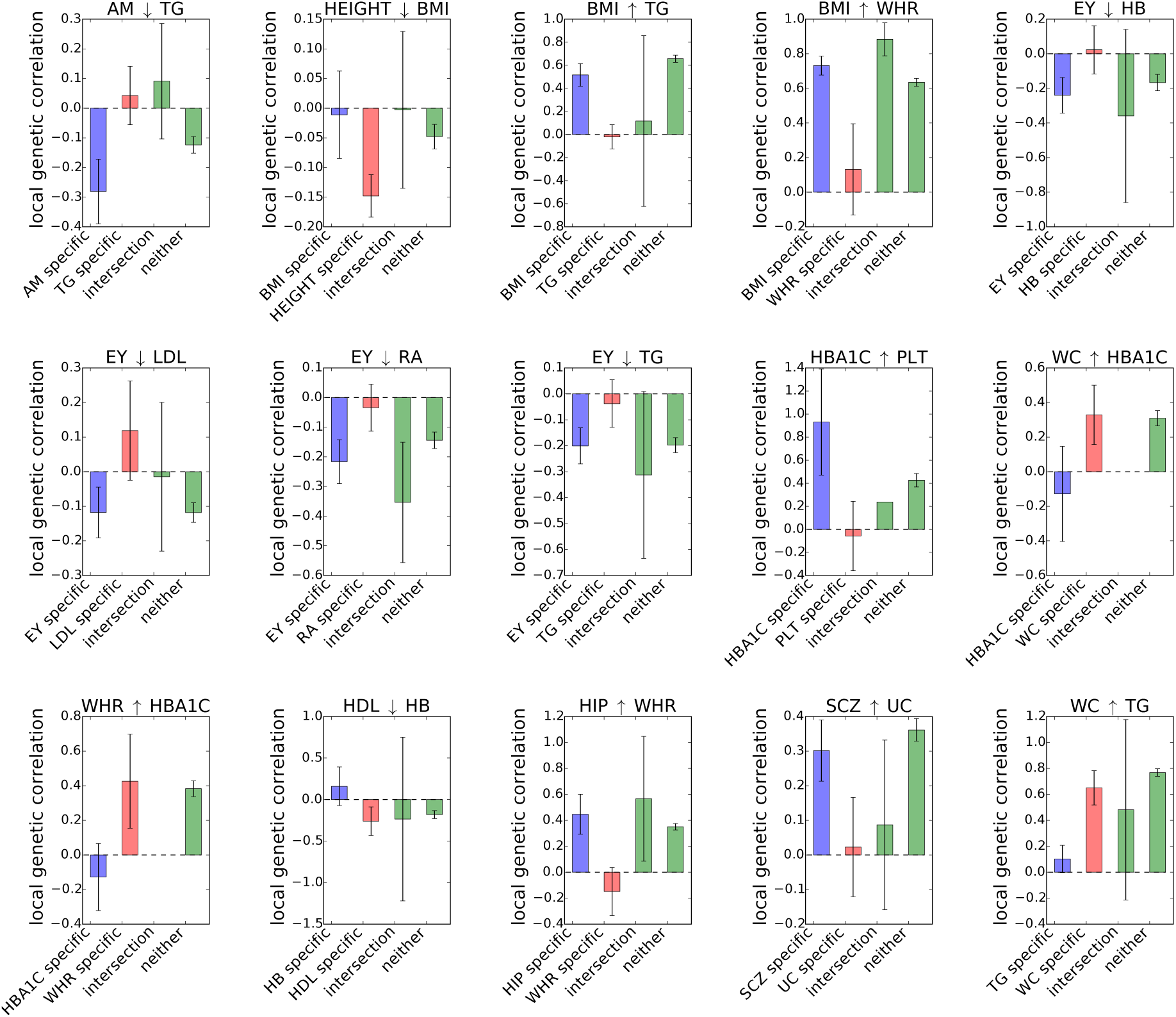
Estimates of local genetic correlation at loci ascertained for GWAS risk variants for each one of 15 pairs of traits that show plausible causal relationship. We obtained standard error using a jackknife approach. Error bars represent 1.96 times the standard error on each side. Here, “↑” means the trait on the left hand side may causally increase the trait on right hand side, whereas “↓”means the trait on the left hand side may causally decrease the trait on the right hand side.

## Discussion

We have described *ρ*-HESS, a method to estimate local genetic covariance and correlation from GWAS summary association data. Through extensive simulations, we demonstrated that our method is approximately unbiased and provides consistent results irrespective of causal architecture. We then applied our method to large-scale GWAS summary association data of 35 complex traits. Compared with cross-trait LDSC, our methods identified considerably more pairs of traits displaying significant genome-wide genetic correlation likely because of the truncated-SVD regularization of the LD matrix, which decreases the standard error of the estimates. We identify genomic regions that are significantly correlated across pairs of traits regardless of the significance of genome-wide correlation. Finally, we performed bi-directional analyses over the local genetic covariances, and report plausible causal relationships for 15 pairs of traits.

We conclude with several limitations of our methods highlighting areas for future work. First, our estimator requires phenotype correlation between two traits, as well as the number of shared individuals between the two GWASs. We estimate the phenotype correlation through cross-trait LDSC assuming full sample overlap between GWAS within the same consortium and no sample overlap between GWAS across two consortium. Although we believe this is supported by the data we analyzed in this study, mis-specifications in the overlapping sample sizes could introduce biases. Second, we emphasize that our bi-directional analyses only identify putative causality and is not proof of causality relations; exact inference of causal relations is complicated by unobserved confounders such as socioeconomic status and/or biological pathways. Furthermore, most of the GWAS summary association data are adjusted for covariates such as age, gender, to increase statistical power^38^, and previous works have shown that adjusting for covariates can potentially lead to false positives^39^. In the bi-directional analyses over local genetic correlation, covariate-adjusted summary association data can potentially lead to false inference of causality. However, the extent to which covariate-adjusted summary association data lead to false discovery of causality remains unclear, and we leave detailed investigation of this issue as future work. Third, in our real data analyses, we made the assumption that the loci are independent of each other. In reality however correlations may exist across adjacent loci due to long range LD, and can lead to biased estimates. Nevertheless, we note that previous works have indicated the effect of LD leakage to be minimal, and we conjecture that this statement still hold in estimating local genetic covariance. Lastly, we uses truncated-SVD to regularize LD matrix and to reduce standard error in the estimates of local genetic covariance, at the cost of introducing bias. Currently, we use a fixed number of eigenvectors in the truncated-SVD regularization, across all the loci. However, this approach may not be optimal for genomic regions with different LD structure, and leave a principled approach of estimating the number of eigenvectors as future work.

## Acknowledgement

We are grateful to Gleb Kichaev, Malika Kumar, Suraj Alva, and James Boocock for their helpful discussions that greatly improved the quality of this manuscript. We also thank Dr. Nicole Soranzo for kindly sharing summary data for the platelet traits. The authors declare no conflict of interest.

## Methods

### Local genetic correlation under fixed-effect model

Let *ϕ*=*x*^T^*β*+*ε* and *ψ*=*x*^T^*γ*+*δ* be two traits measured at an individual, standardized so that E[*ϕ*] = E[*ψ*] = 0 and Var[*ϕ*] = Var[*ψ*] = 1, where *β*, *γ*∊ℝ*^p^* are the fixed effect size vectors for the two traits; *x*∊ℝ*^p^*, the genotype vector of the individual at *p* SNPs, standardized so that E[*x*] = 0, and Var[*x*] = ***V***, the LD matrix; and ε,δ, environmental effects independent of *x*, *β*, *γ*, with E[*ε*] = E[*δ*] = 0, Var[*ε*] = *σ*^2^*_ε_*, Var[*δ*] = *σ*^2^*_δ_*, and Cov[*γ*, *δ*] = *ρ_e_*. Under these assumptions, one can decompose the phenotypic covariance, *ρ*, between *ϕ* and *ψ* as

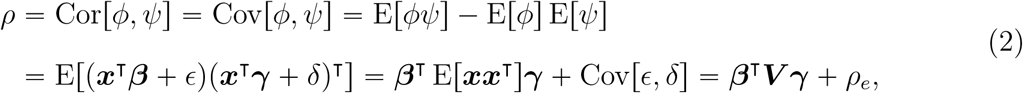

where *ρ_g_*=*β*^T^***V****γ* is the genetic covariance between the two traits. For standardized traits, phenotypic covariance coincides with phenotypic correlation. Thus, given the true effect size vectors, *β*, *γ*, and the LD matrix ***V***, one can easily obtain *ρ_g_*. We define genetic correlation, *r_g_*, as 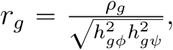, i.e. genetic covariance standardized by square root of SNP-heritability of the two traits.

### Estimating local genetic covariance from GWAS summary data

In two GWASs involving *n_1_* individuals for trait 1 (*ϕ*), *n_2_* individuals for trait 2 (*ψ*), and *n_s_* shared individuals, we assume

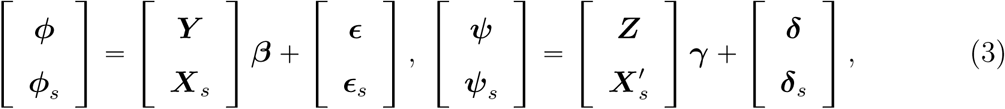

where (*ϕ*, *ϕ_S_*)∊ℝ^*n*1^and (*ψ*.*ψ_S_*)∊ℝ^*n*2^ are the standardized trait values of all individuals in each GWAS; (***Y***, ***X****_s_*)∊ℝ^*n*1×*p*^, (***Z***, ***X′****_s_*)∊ℝ^*n*2×*p*^, column standardized genotype matrices of all individuals in each GWAS, where ***X****_S_* and ***X′****_S_* represent the genotype matrices for the same set of individuals and SNPs but standardized differently in each GWAS; (*ε*, *ε_S_*)∊ℝ^*n*1^,(*δ*, *δ_S_*)∊ℝ^*n*2^, environmental effects of all individuals in each GWAS. We use the subscript ‘s’ to represent individuals shared by both GWASs. We further assume that E[*ε*] = E[*δ*] =0 E[*ε_S_*] = E[*δ_S_*] = 0, Var[*δ*] = Var[*δ_S_*] = *σ*^2^*_δ_****I***, Var[*δ*] = Var[*δ_s_*] =*σ*^2^*_δ_****I***, Cov[*ε*, *δ*] = 0, and Cov[*ε_S_*, *δ_S_*] = *ρ*_*e*_***I***.

In a traditional GWAS, we obtain marginal effect size estimates, 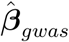 and 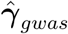, as

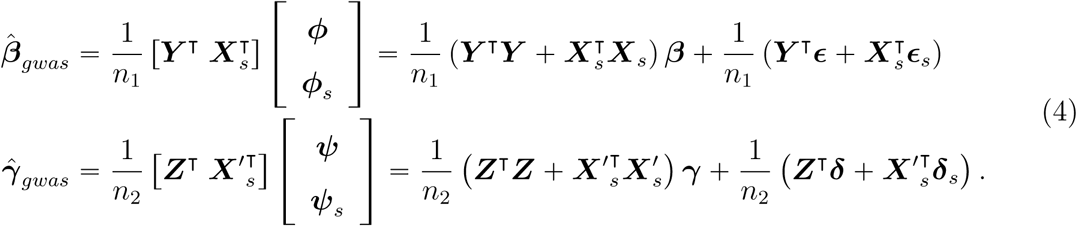

Assuming individuals in both GWASs are drawn from the same population with LD matrix ***V***, we have 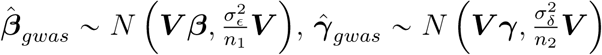 we also find

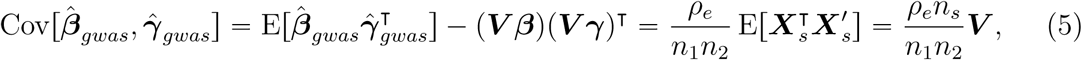

where the last equality follows from Isserlis’ theorem^40^.

Under infinite sample sizes, 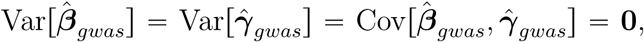 and we 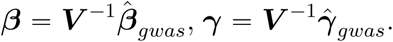Thus, local genetic covariance, *ρ*_*g,local*_, can be computed as

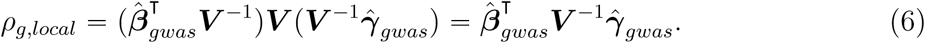

However, when sample sizes are finite, from bilinear form theory^41^, the covariance between 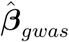 and 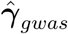 creates bias, resulting in

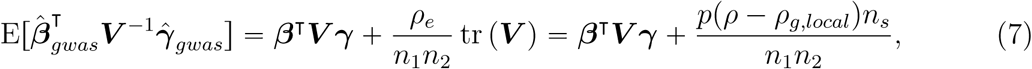

Correcting for bias, we arrive at the unbiased estimator

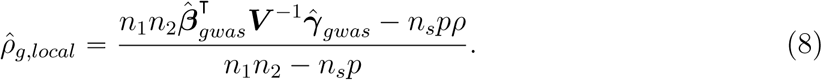

For rank-deficient LD matrix ***V***, one replaces ***V*** ^−1^ with the pseudo-inverse (***V***^†^) and *p* with *q* = rank(***V***), yielding the unbiased estimator

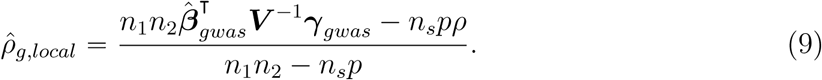

Thus, in order to obtain an unbiased estimate of genetic covariance between a pair of traits, one needs to know their phenotypic correlation. When phenotypic correlation is not available, one can obtain an estimate from genome-wide summary data using the LD score regression equation^14^,

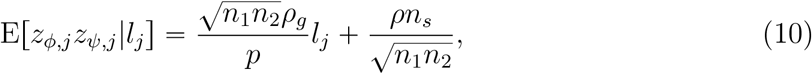

where *z_ϕ,j_*, *z_ψ,j_* are the Z-scores of SNP *j* in the two traits, and *l_j_* the LD score of SNP *j*.

In the special case when 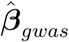 and 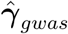 are obtained for the same trait on the same set of individuals (i.e. 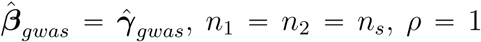) Equation (8) reduces to the local SNP-heritability estimator^18^. When *n_s_* = 0 (i.e. no shared individuals between the GWASs), the unbiased estimator is simply 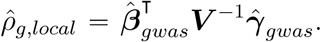 An interpretation for this simple formula is that in the absence of sample overlap, the covariance in the noise, *ε* and *δ*, is 0 and does thus not introduce bias into the estimate of *ρ*_*g,local*_.

Following bilinear form theory^41^, we obtain the variance of 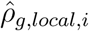,

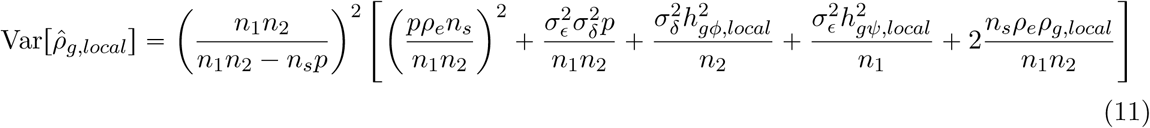

For rank deficient LD matrix with rank(***V***) = *q*, one replaces *p* with *q* in Equation (11).

### Accounting for statistical noise in LD estimates

Limited sample size of external reference panels creates statistical noise in the estimated LD matrix that biases our estimates. Following our previous work^18^, we apply truncated-SVD regularization to remove noise in external reference LD. We note that 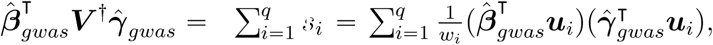, where *w_i_*, ***u**_i_* are the eigenvalues and eigenvectors of the LD matrix ***V***, and *q* = rank(***V***). We use 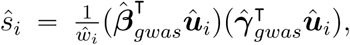 to denote the counterpart obtained from external reference LD matrix ***V̂***. We show through simulations that the bulk of 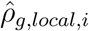 comes from *s_i_* where *i* ≪ *q* and that *s_i_* ≈ *ŝ_i_* for *i* ≪ *q*, thus justifying truncated-SVD as an appropriate regularization method when only external reference LD (***V̂***) is available.

Let 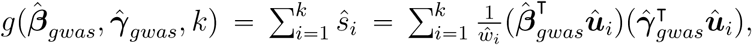 be the truncated-SVD regularized estimates for 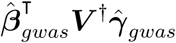 then it can be shown that

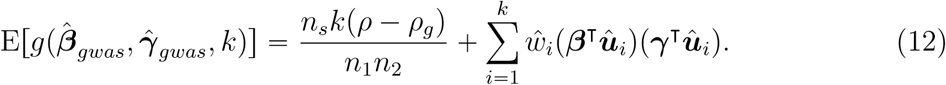

Assuming *Ŵ_i_* = *w_i_* and *û_i_* = *u_i_* for *i* ≪ *k*, Equation (12) is a biased approximation of *ρ*_*g,local*_, with bias 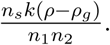 Correcting for the bias, we arrive at the estimator

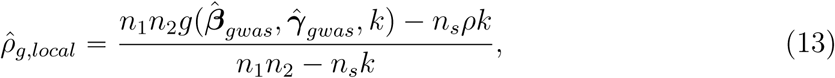

which has variance

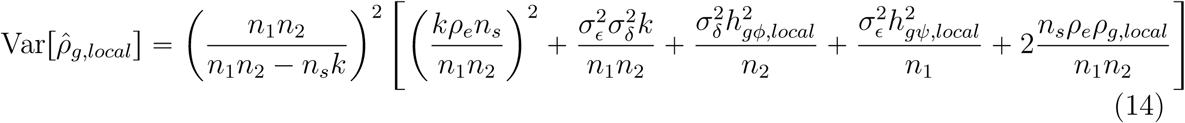

### Extension to multiple independent loci

For genome partitioned into *m* loci, let

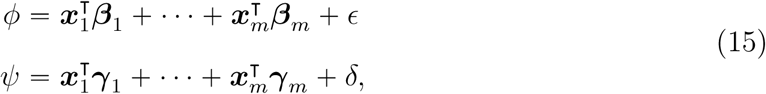

denote the phenotype measurements of two traits at an individuals, where we assume that SNPs in different pairs of loci are independent, i.e. E[*x_ik_x_il_*] = 0 for all *i*≠*j*, *k* ∊ {1, ⋯, *p_i_*}, and *l* ∊ {1, ⋯, *p_j_*}, where *p_i_* and *p_j_* are the number of SNPs in locus *i* and *j*. Under these assumptions, we decompose the phenotypic covariance, *ρ*, between *ϕ* and *ψ*,

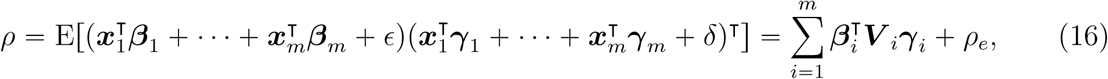

where 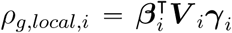 is the genetic covariance between the pair of traits attributed to genetic variants at locus *i*. Following strategies outlined in previous sections, we arrive at the estimator for genetic covariance at the *i*-th locus,

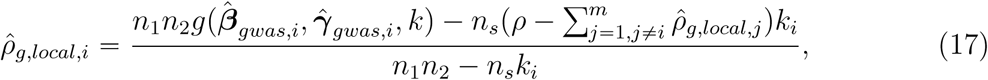

which defines a system of linear equation involving *m* unknown variables and *m* equations. In the special case where there is no sample overlap, *n_s_* = 0, and 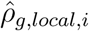 reduces to 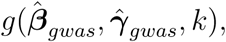 and can be estimated independent of all other windows. Following bilinear form theory, we obtain variance estimate for 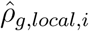 as,

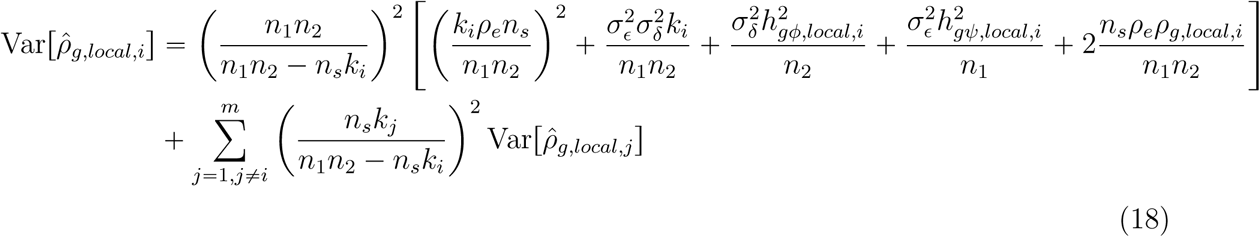

which also defines a system of linear equations with *m* equations and *m* variables.

When *k*_1_ = ⋯ = *k_m_* = *k*, i.e. all loci use the same number of eigenvectors in the truncated-SVD regularization, summing over *i* on both sides of Equation (17) yields

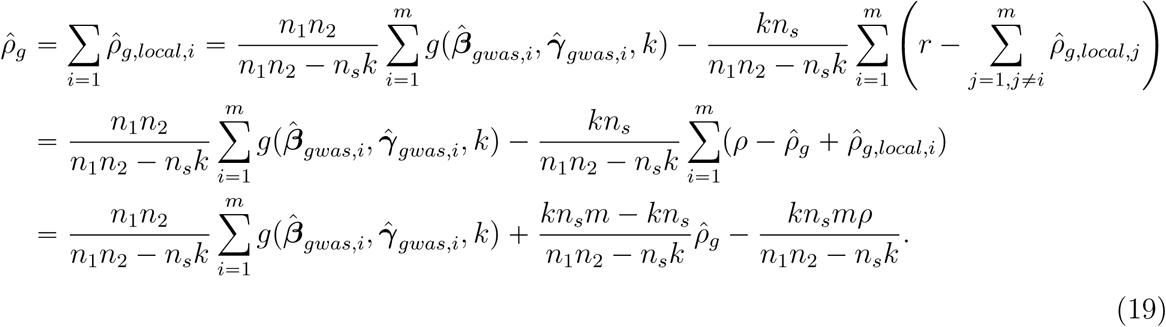

Solving for *ρ̂_g_* yields

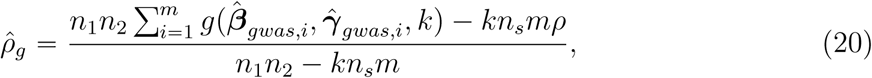

which has variance

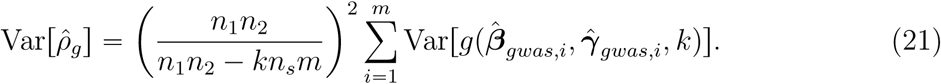

Thus, if *k* is chosen such that (*n*_1_*n*_2_−*kn_s_m*) is small (i.e.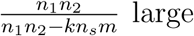 large), the estimate of total genetic covariance will have large standard error. To reduce standard error in the estimates (at the cost of some bias), we recommend choosing *k* such that 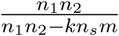 is less than 2. When testing for statistical significance, we assume that the estimates of local and genome-wide genetic covariance and correlation follow a normal distribution.

### Standardizing local genetic covariance

Standardizing local genetic covariance between a pair of traits by the square roots of the local heritability of the two traits yields the local genetic correlation (*r_g,local_*). Whereas genetic covariance takes into account the magnitude of the effect sizes, genetic correlation provides a measure of similarity between the effects of SNPs on traits comparable across different magnitudes of effect sizes. To estimate local genetic correlation for the *i*-th locus, we apply the formula

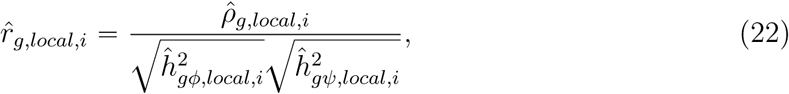

where 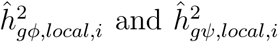 denote the local SNP-heritability of trait *ϕ* and *ψ* at the *i*-th locus. In simulations, we show that 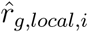 is approximately unbiased when both traits are heritable at the *i*-th locus. In practice, however, the terms, 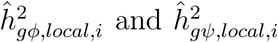 can be close to zero, greatly inflating the standard error of 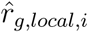 Thus, we recommend estimating local genetic correlation only at loci with significant local SNP-heritability. One can also estimate local genetic correlation at a set of loci. For example, to estimate genetic correlation at loci indexed by the index set *C*, one applies the following formula,

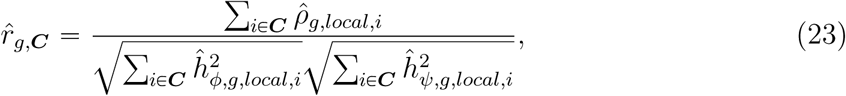

We estimate standard error of local genetic correlation at a single locus through a parametric bootstrap approach^42^ and local genetic correlation at a set of loci through jackknife.

### Simulation framework

Starting from half (202 individuals) of the EUR reference panel from the 1000 Genomes Project^22^, we simulated genotype data for 50,000 individuals at HapMap3^43^ SNPs with minor allele frequency (MAF) greater than 5% in a 1.5Mb locus on chromosome 1 using HAPGEN2^43^. We used the other half of the EUR reference panel (203 individuals) to obtain external reference LD matrices.

We simulated phenotypes from the genotypes according to the linear model *ϕ=****X****β*+*ε* and *ψ*=***X****γ* +,***δ***, where ***X*** is the column-standardized genotype matrix. We drew the effects of causal SNPs (*β*_*C*_, *γ*_*C*_) from the distribution

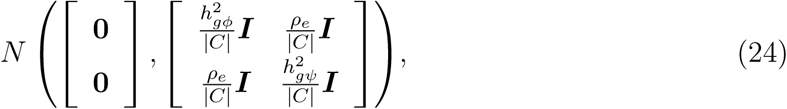

where *C* is the index set of causal SNPs, and set the effects of all other SNPs to be zero. Here, for the convenience of simulation, we assume both traits have the same causal SNPs, although this assumption doesn’t need to hold for *ρ*-HESS to be unbiased. We then drew (***ε***, ***δ***) from the distribution

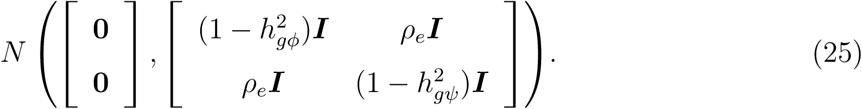

Finally, we simulated GWAS summary statistics using methods outlined in previous sections. For each ***β*** and *γ* drawn from the normal distribution, we simulated 500 sets of summary statistics by varying ***ε*** and ***δ***, and applied *ρ*-HESS to estimate genetic covariance and genetic correlation for each set of the simulated summary statistics.

### Inferring direction of causality

We systematically search for plausible causal directions (see example 1 and 2 in Figure 9) across all 234 pair of traits that show significant genome-wide correlation. First, we test whether the 95% confidence intervals, defined as 1.96 times standard error on each side of the estimates, of 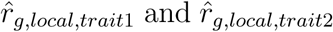 overlap. Then we test whether one of the confidence intervals overlaps with 0 and the other does not. Since we employed jackknife to estimate standard errors, for the robustness of the standard error, we restricted our bi-directional analyses to pairs of traits for which the number of loci ascertained for GWAS risk variants specific to both traits is greater than 10. We mark pairs of trait, for which the confidence intervals of local genetic correlations do not overlap, as pairs of traits having putative causal relationships, and designate the trait, for which the local genetic correlation is significantly non-zero, as the causal trait.

### Empirical data sets

We obtained GWAS summary data for 35 complex traits and diseases from 11 GWAS consortia (see Table 1), all of which are based on individuals of European ancestry, and have sample size greater than 20,000. We used approximately independent loci defined in^21^ to partition the genome, and restricted our analyses on HapMap3 SNPs with minor allele frequency (MAF) greater than 5% in the European population in the 1000 Genomes data^22^. We also removed stand-ambiguous SNPs prior to our analyses. We follow the method outlined in^18^ to estimate and re-inflate λ _*gc*_, and to choose the number of eigenvectors to include in estimating local genetic covariance and SNP-heritability. We note that although adjustment factors are needed to correct for ascertainment bias in the estimation of local SNP-heritability and local genetic covariance^18^, they are not needed in the estimation of local genetic correlation^14^.

**Table 1:** A summary of the 35 GWAS summary data sets analyzed. ^a^Total number of traits with significant non-zero genome-wide genetic correlation (two-tailed *p* < 0.05/595). ^b^Total number of traits outside the consortium with significant non-zero genome-wide genetic correlation. Number of traits for which the magnitude of genetic correlation is both significantly non-zero and greater than 0.2 is shown in parentheses.

| Trait Name | Abbreviation | Consortium | # gen corr <sup>a</sup><br>within<br>consortium | # gen corr <sup>b</sup><br>outside<br>consortium | Approx.<br>sample size |
| --- | --- | --- | --- | --- | --- |
| Body Mass Index<sup>44</sup> | BMI | GIANT | 23 (16) | 19 (13) | 231K |
| Height<sup>45</sup> | HEIGHT | GIANT | 17 (2) | 13 (0) | 241K |
| Hip Circumference<sup>46</sup> | HIP | GIANT | 21 (14) | 17 (10) | 144K |
| Waist-to-hip Ratio<sup>46</sup> | WHR | GIANT | 22 (16) | 18 (13) | 143K |
| Waist Circumference<sup>46</sup> | WC | GIANT | 23 (17) | 19 (13) | 153K |
| Haemoglobin<sup>30</sup> | HB | HaemGen | 15 (9) | 12 (8) | 51K |
| Mean Cell Haemoglobin<sup>30</sup> | MCH | HaemGen | 7 (1) | 6 (1) | 44K |
| MCH Concentration<sup>30</sup> | MCHC | HaemGen | 6 (4) | 1 (1) | 47K |
| Mean Cell Volume<sup>30</sup> | MCV | HaemGen | 10 (3) | 8 (1) | 49K |
| Packed Cell Volume<sup>30</sup> | PCV | HaemGen | 15 (10) | 11 (8) | 45K |
| Red Blood Cell Count<sup>30</sup> | RBC | HaemGen | 17 (10) | 14 (8) | 46K |
| Number of Platelets<sup>47</sup> | PLT | HaemGen | 10 (1) | 6 (1) | 67K |
| Fasting Glucose<sup>48</sup> | FG | MAGIC | 18 (10) | 15 (9) | 46K |
| Fasting Insulin<sup>48</sup> | FI | MAGIC | 18 (12) | 16 (12) | 46K |
| HbA1C<sup>49</sup> | HbA1C | MAGIC | 18 (14) | 16 (13) | 46K |
| HOMA-B<sup>48</sup> | HOMA-B | MAGIC | 16 (9) | 13 (9) | 46K |
| HOMA-IR<sup>48</sup> | HOMA-IR | MAGIC | 16 (13) | 16 (13) | 46K |
| High Density Lipoprotein<sup>26</sup> | HDL | GLGC | 18 (11) | 16 (10) | 96K |
| Low Density Lipoprotein<sup>26</sup> | LDL | GLGC | 15 (6) | 13 (4) | 91K |
| Total Cholesterol<sup>26</sup> | TC | GLGC | 15 (4) | 12 (2) | 96K |
| Triglycerides<sup>26</sup> | TG | GLGC | 22 (13) | 19 (10) | 92K |
| Education Years<sup>50</sup> | EY | SSGAC | 25 (8) | 22 (6) | 294K |
| Depressive Symptoms<sup>51</sup> | DS | SSGAC | 10 (4) | 7 (1) | 161K |
| Neuroticism<sup>51</sup> | NEURO | SSGAC | 5 (3) | 2 (0) | 171K |
| Subjective Well-being<sup>51</sup> | SWB | SSGAC | 7 (2) | 4 (0) | 298K |
| Forearm BMD<sup>52</sup> | FA | GEFOS | 3 (1) | 1 (0) | 53K |
| Femoral Neck BMD<sup>52</sup> | FN | GEFOS | 4 (2) | 2 (0) | 53K |
| Lumbar Spine BMD<sup>52</sup> | LS | GEFOS | 5 (1) | 3 (0) | 53K |
| Age at Menarche<sup>53</sup> | AM | ReproGen | 17 (3) | 17 (3) | 133K |
| Rheumatoid Arthritis<sup>54</sup> | RA | RACI and GARNET | 3 (0) | 3 (0) | 58K |
| Schizophrenia<sup>55</sup> | SCZ | PGC | 13 (4) | 13 (4) | 75K |
| Crohn's Disease<sup>56</sup> | CD | IIBD | 5 (4) | 3 (2) | 52K |
| Inflammatory Bowel Disease<sup>56</sup> | IBD | IIBD | 5 (4) | 3 (2) | 66K |
| Ulcerative Colitis<sup>56</sup> | UC | IIBD | 4 (4) | 2 (2) | 48K |
| Type 2 Diabetes<sup>57</sup> | T2D | DIAGRAM | 20 (13) | 20 (13) | 62K |

